# Dynamic Gating by the Phenylalanine Clamp Loop Controls Peptide Translocation Through the Anthrax Toxin Nanopore

**DOI:** 10.1101/2025.09.25.678637

**Authors:** Jennifer M. Colby, Bryan A. Krantz

## Abstract

The ϕ-clamp loop, which contains the key F427 residue, is a critical active site in the anthrax toxin protective antigen (PA) nanopore, yet its precise role in governing the complex, multi-state dynamics of peptide translocation remains debated. Here, we dissect the peptide-clamp interaction mechanism using single-channel electrophysiology and a series of guest-host peptides, which are translocated via either wild-type PA, an ablated F427A mutant, or a polar aromatic F427Y mutant. Mutations to the ϕ clamp dramatically reduce peptide residence times but, critically, preserve the intermediate, partially blocked conductance states observed in the wild-type pore. Thermodynamic analysis reveals that the F427 residue is essential for creating a deep, energetically stable, fully blocked ‘hydrophobic trap’ (State 0), as its mutation leads to a significant destabilization of this state and a corresponding population shift to shallower intermediates. Kinetic analysis of the state-to-state transitions demonstrates that while the F427A mutation lowers the energetic barrier for escape from this trap, it disrupts the efficient, hydrophobically driven entry. Furthermore, the strong correlations between kinetic parameters and peptide molecular properties (hydrophobicity, aromaticity) that are a hallmark of the wild-type pore are completely abolished in the F427A mutant. These results support a refined model where F427 acts as a specific chemical ‘reader,’ and the intermediate states arise from larger-scale, dynamic dilation of the entire clamp-containing loop. This detailed mechanistic insight provides a framework for the rational engineering of next-generation nanopore biosensors.

## Introduction

The anthrax toxin protective antigen (PA) nanopore has emerged as a premier model system for investigating the biophysics of protein translocation (1). The synergy between high-resolution structural biology (2–5) and dynamic single-channel analysis has enabled the development of detailed mechanistic models for its function. This natural translocase is exceptionally robust, processively transporting peptides under a voltage or proton gradient (6, 7) with high sensitivity, capable of detecting unlabeled substrates at nanomolar concentrations (8–14). Central to its function are internal ‘peptide-clamp’ sites, most notably the phenylalanine clamp (ϕ clamp), a ring of F427 residues that forms non-specific yet critical interactions with the substrate (3, 4, 9, 11, 15, 16). These dynamic interactions produce complex, multi-state ionic current signatures, each with distinct kinetic and conductance characteristics (9, 11–13). While these signatures are information-rich, deciphering them to map the translocation energy landscape and assign physical drivers to each kinetic step remains a formidable challenge. The information-rich nature of these signals has also spurred the development of the PA nanopore as a powerful biosensor, particularly when combined with machine learning for high-resolution analyte classification (12, 13). Further advances in this area hold promise for the long-term goal of single-molecule peptide sequencing, a rapidly emerging field of research (17–19).

A deeper understanding of the dynamics governing peptide-clamp interactions is therefore fundamental, serving as a linchpin for both advancing the basic science of translocation and enabling the translational development of next-generation biosensors and sequencers. These dynamics, potentially arising from the population of alternate ϕ-clamp conformations, are rich with mechanistic and analytical information. Consequently, systematically modifying these interactions through rational nanopore engineering is a promising strategy for creating biosensors with enhanced capabilities for detecting challenging analytes (13).

Given that aromatic, clamp-like structures are a recurring motif in many biological translocases and unfoldases, a detailed investigation of the PA ϕ-clamp offers broadly applicable insights. Here, we dissect the role of the F427 residue through targeted mutagenesis (F427A and F427Y), moving beyond qualitative observation to a quantitative thermodynamic and kinetic deconstruction of the translocation mechanism. We map the energy landscape of the peptide-clamp interaction, defining the stability of a deeply bound ‘hydrophobic trap’ and quantifying the energetic barriers for transitions between it and other intermediate states. Furthermore, our results provide strong evidence for a refined structural model, suggesting that the intermediate conductance states arise not from simple side-chain rotamers but from a larger-scale, dynamic dilation of the entire ϕ-clamp-containing loop.

## Results

In this PA nanopore system, the ϕ clamp forms a structural bottleneck in all observed cryo-EM structures (**Fig. 1A**). To probe the role of the F427 residue in the ϕ clamp in the observed multi-state peptide translocation mechanism, two F427X mutants were investigated: (i) F427A, which completely ablates the aromatic and hydrophobic side chain; and (ii) F427Y, which substitutes it with a polar yet aromatic residue. A 10-residue guest-host peptide of sequence KKKKKXXSXX with various guest site, X, chemistries (abbreviated throughout by their three-letter notation) included Ala, Leu, Phe, Thr, Trp, and Tyr. One peptide (called ‘guest-host TrpDL’), where the guest residue was Trp but every other residue was ᴅ chirality, was distinct conformationally from guest-host Trp, which, like the other peptides, was comprised of uniform natural ʟ stereoisomers. This peptide series produced rich dynamic translocation kinetics with the wild-type ϕ clamp, hence we again employed these peptides to investigate the thermodynamics and kinetics in the context of ϕ clamp variants. The most striking qualitative observation from the raw data (**Figs. 1B, S1)** is that mutations to F427 lead to a dramatic reduction in the overall residence time of the peptides. The effect is most pronounced for the F427A mutant, suggesting the F427 residue is a primary determinant of peptide trapping.

**Fig. 1.**
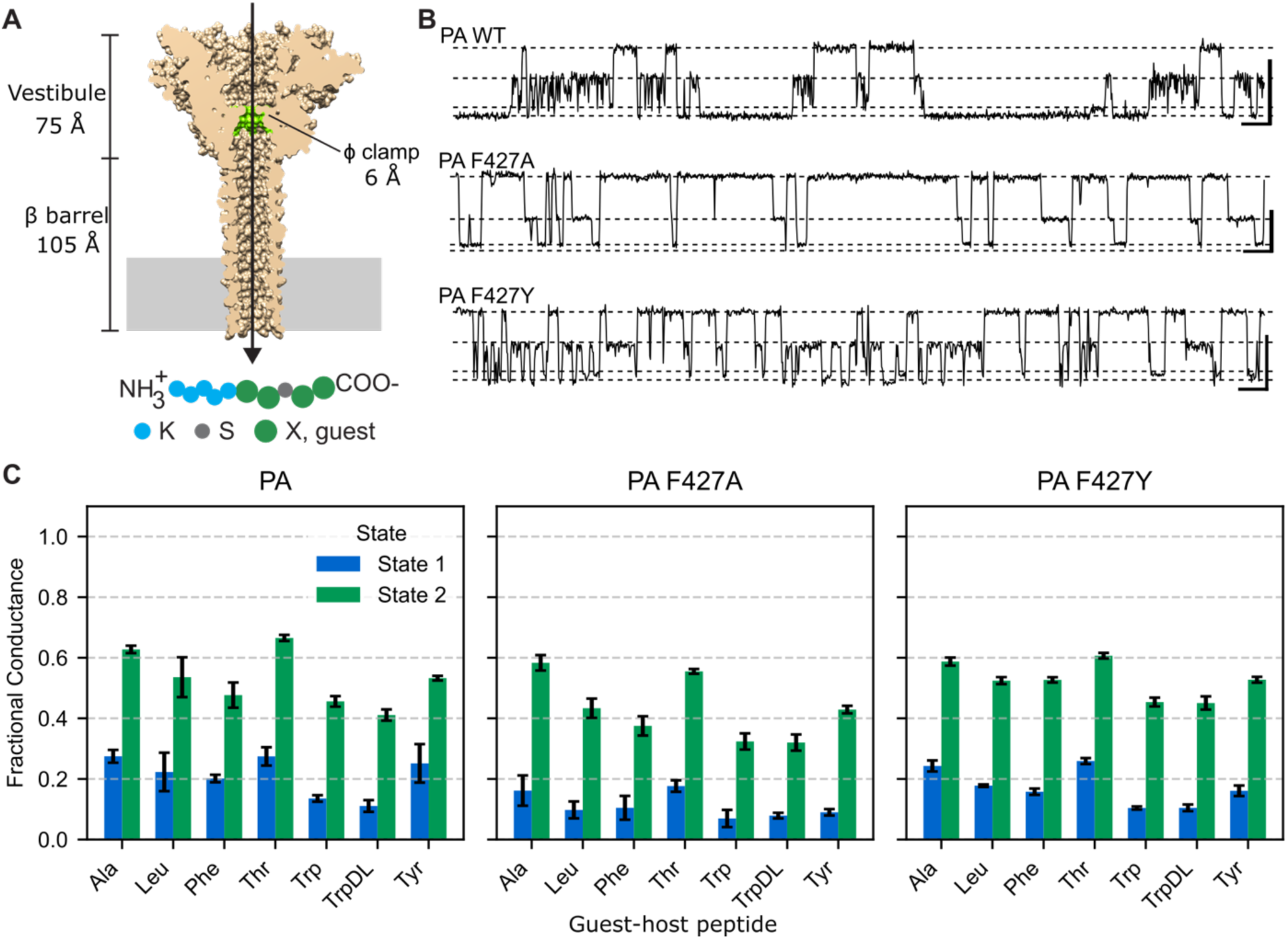
Single-channel analysis of guest-host peptide translocation via ϕ-clamp variants of PA nanopore. **(A)** (upper) Sagittal section of molecular surface of PA nanopore highlighting the F427 ϕ-clamp loop (green). (lower) Schematic of the 10-residue guest-host peptide of sequence, KKKKKXXSXX, where guest residue is X. **(B)** Selected raw translocation event streams of guest-host Tyr peptides via WT PA, PA F427A, and PA F427Y nanopores at 70 mV in 100 mM KCl, pH 5.6. Discrete conductance levels for states populated during translocation are indicated by dotted lines, where the topmost of each panel is the open pore (State 3) and the bottommost is the fully blocked pore (State 0); the intermediate partially blocked species (State 1 and State 2) are in between. The scalebars to the right for each nanopore are 4 pA by 100 ms. Note PA F427A conducts more than PA F427Y and WT PA. (See also **Table S1** for the absolute conductance of open pore for these respective nanopores). The full set of slices of peptide translocation event streams described in this study are plotted in **Fig. S1**. **(C)** Comparison of fractional conductance levels of State 1 and State 2, which have intermediate fractional conductance for the WT PA (left), PA F427A (middle), and PA F427Y (right) nanopore. Fractional values were estimated for each guest-host peptide under the conditions described in panel B.

### Persistence of intermediate states in F427X mutant nanopores

Despite the observed kinetic changes in peptide translocation for the mutant nanopores, the four-state conductance model, including the distinct partially conducting intermediate states (State 1 and State 2), is essentially conserved across all three nanopore variants (**Fig. 1C, Supporting Document 1)**. There are some second order trends in the fractional conductance levels of the State 1 and State 2 intermediates, where bulkier peptides show slightly deeper blockade depths; however, consistently the State 2 species is ∼50% max conducting. Overall, these data argue that the physical basis for these conductance state intermediates is not simply due to the population of alternate F427 side chain rotamer, but rather it is likely there is a larger-scale conformational change of the entire ϕ clamp-containing loop—a persistent feature that is preserved even in the mutants.

### ϕ clamp forms an electrophysiological bottleneck in the pore

Both the F427A and F427Y mutations lead to a statistically significant increase in the open pore State 3 conductance (108 and 80 pS, respectively) relative to wild type (68 pS) **(Table S1)**. These measurements at pH 5.6 replicate earlier work at pH 6.6 (15), further solidifying that the F427 residue electrophysiologically forms the narrowest, most hydrophobic constriction or bottleneck for ion flow in the wild-type nanopore. For the observed peptide-clamp species, we have proposed the dynamic and partially conducting states observed across peptide classes (**Fig. 1C**) are putative peptide-liganded but dilated conformers of this bottleneck site (14).

### Thermodynamic analysis shows ϕ-clamp creates deep hydrophobic trap

To understand how the mutations alter the energy landscape of the observed peptide complexes during translocation, the relative free energy of each bound state was calculated from the fractional state occupancies or probabilities. The free energy change (ΔΔ*G*) for each F427X variant and peptide combination was determined using the wild-type PA nanopore as the reference state. The most significant finding from the thermodynamic analysis is that the F427A mutation consistently and dramatically destabilizes State 0 for all peptides, as shown by the large, positive ΔΔ*G* values (**Fig 2, Supporting Document 2)**. The F427Y mutant has a similar but less pronounced effect, demonstrating that a ϕ clamp with more hydrophobic character creates a deeper energy well. Therefore, the wild-type F427 residue comprises a key structural element responsible for creating the deep, stable ‘hydrophobic trap’ of the fully blocked State 0. The consequence of destabilizing State 0 is accompanied by a relative stabilization (negative ΔΔG) of States 1 and 2, particularly for the more strongly interacting peptides. This effect is a classic thermodynamic population shift, where making the deepest energy well shallower with the F427X mutations cause the peptide population to redistribute into the intermediate, partially blocked conductance states.

**Fig. 2.**
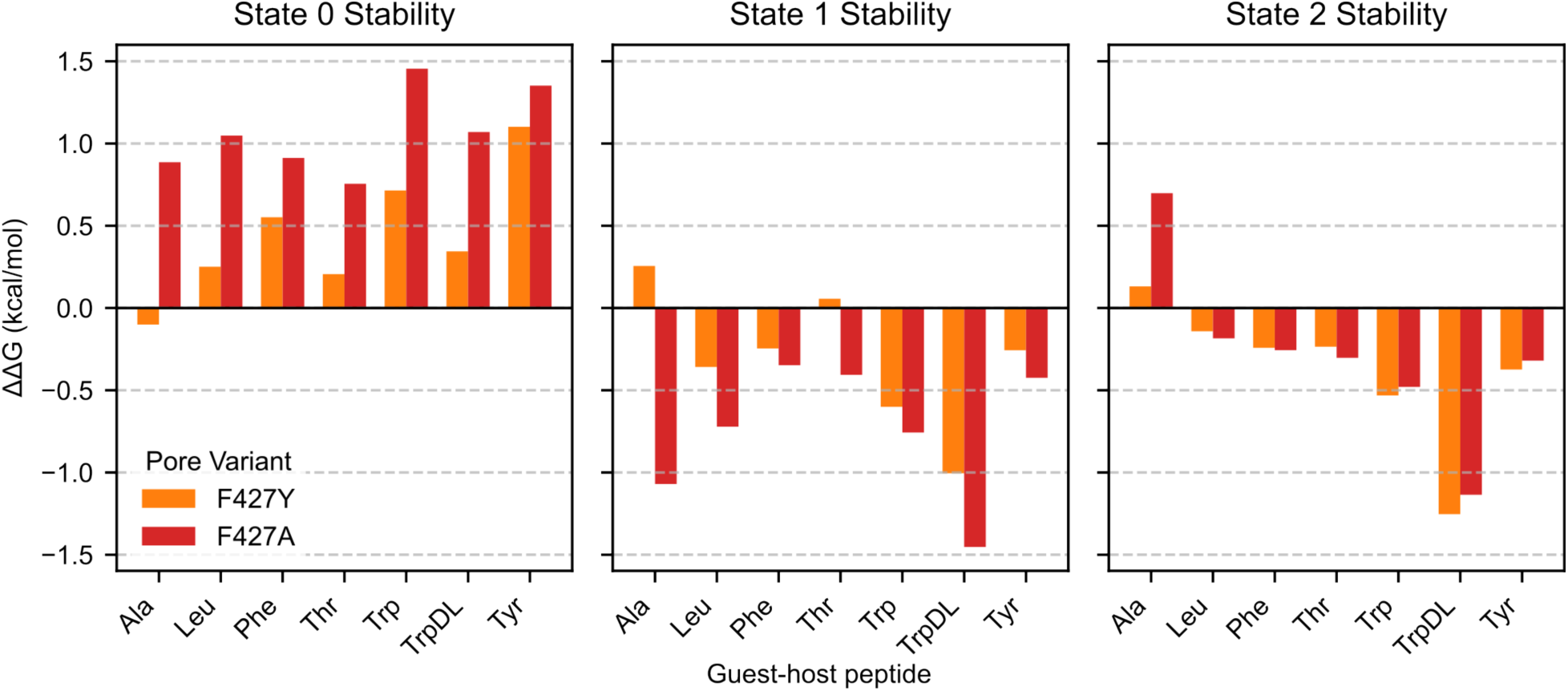
Mutations to the ϕ-clamp alter the thermodynamic stability of peptide-bound states. The change in thermodynamic stability (ΔΔG) for the three bound conductance states— State 0 (left), State 1 (middle), and State 2 (right)—caused by the F427A (red) and F427Y (orange) mutations. ΔΔG was calculated from the fractional state occupancies derived from aggregated single-channel dwell times. Positive values indicate the mutation destabilizes a given state (making it less populated), while negative values indicate stabilization relative to the wild-type pore. Note the consistent and significant destabilization of the deeply bound State 0 by the F427A mutation across all peptides, which is accompanied by a corresponding stabilization of the intermediate states.

### Kinetic analysis identifies role of F427 in controlling translocation barriers

A 4×4 dwell time transition matrix was determined for each guest-host peptide from the complete set of translocation events for each F427X nanopore variant. Survival curves were generated from the dwell times for each transition represented in the dataset. To compute lifetimes (τ) and amplitudes (*A*), one-, two- and three-exponential decays were fitted to the natural log of the survival coordinate. Many transitions for these peptides’ translocations deviated from linearity and exhibited two- or three-exponential relationships, indicating the presence of hidden states sharing the same conductance levels **(Fig. S2)**. These results for the F427X nanopore variants are consistent with prior data, where longer peptides (9, 10) and these guest-host peptides (14) translocated through wild-type PA nanopore via hidden states.

To optimize exponential decay model selection for each transition, peptide, and nanopore variant, the Bayesian Information Criterion was used. A mean lifetime τ value (τ_mean_) was computed to aid in comparison of peptides when different exponential decay models were employed for a given transition. To facilitate analysis, τ values from different decay models were handled systematically by ranking as τ_fast_, τ_middle_, and τ_slow_. When the model was a one-exponential decay, then only τ_fast_ was used. When a two-exponential model was used τ_fast_ was the fastest decay, and τ_middle_ and τ_slow_ were the same second slower decay rate. The respective amplitude values were handled similarly to the τ values in this kinetic parameter consolidation process **(Supporting Document 3)**.

Estimates of energy barrier heights between the observed transitions (Δ*G*‡) were determined from τ_mean_ values by *RT* ln τ_mean_. The change in the activation free energy (ΔΔ*G*‡) was calculated to quantify how the F427X mutations affect the energy barriers for specific transitions between states: ΔΔ*G*‡ = Δ*G*‡(F427X) - Δ*G*‡(WT). Hence negative ΔΔ*G*‡ indicate that the mutation lowers the activation energy barrier for a given transition, making it faster than in the wild-type pore; and positive ΔΔ*G*‡ indicates the mutation raises the activation energy barrier, making the transition slower than in the wild type. The complete analysis of ΔΔ*G*‡ values for all transitions **(Supporting Document 4)** is shown **(Fig. S3)**.

Three key steps in mechanism illustrate how changes in ΔΔ*G*‡ propagate consistently across the panel of peptides (**Fig. 3**). From the preceding thermodynamic analysis State 0 represents the most deeply bound species, which in the wild-type context is consistent with a hydrophobic trap that is destabilized by the F427A mutation. Kinetically, the barrier for escaping the trap (State 0→1) is dramatic lowered by the F427X mutations, especially for aromatic peptides with F427A. Entering the hydrophobic trap (e.g., State 1→0) is likewise affected, where F427X mutations can raise this barrier. In the dissociation step (State 1→3), where the peptide leaves the channel, the F427A mutation significantly lowers the barrier for escape, thereby confirming the ‘sticky’ nature of the wild-type ϕ clamp. From this kinetic analysis we conclude that the F427 residue plays a complex, state-dependent role. An intact ϕ-clamp site in the nanopore not only facilitates entry into the most trapped state but also creates a high barrier to escape. Furthermore, it significantly increases the energy barrier for final dissociation, thereby acting as the primary determinant of the long residence times observed in the wild-type nanopore.

**Fig. 3.**
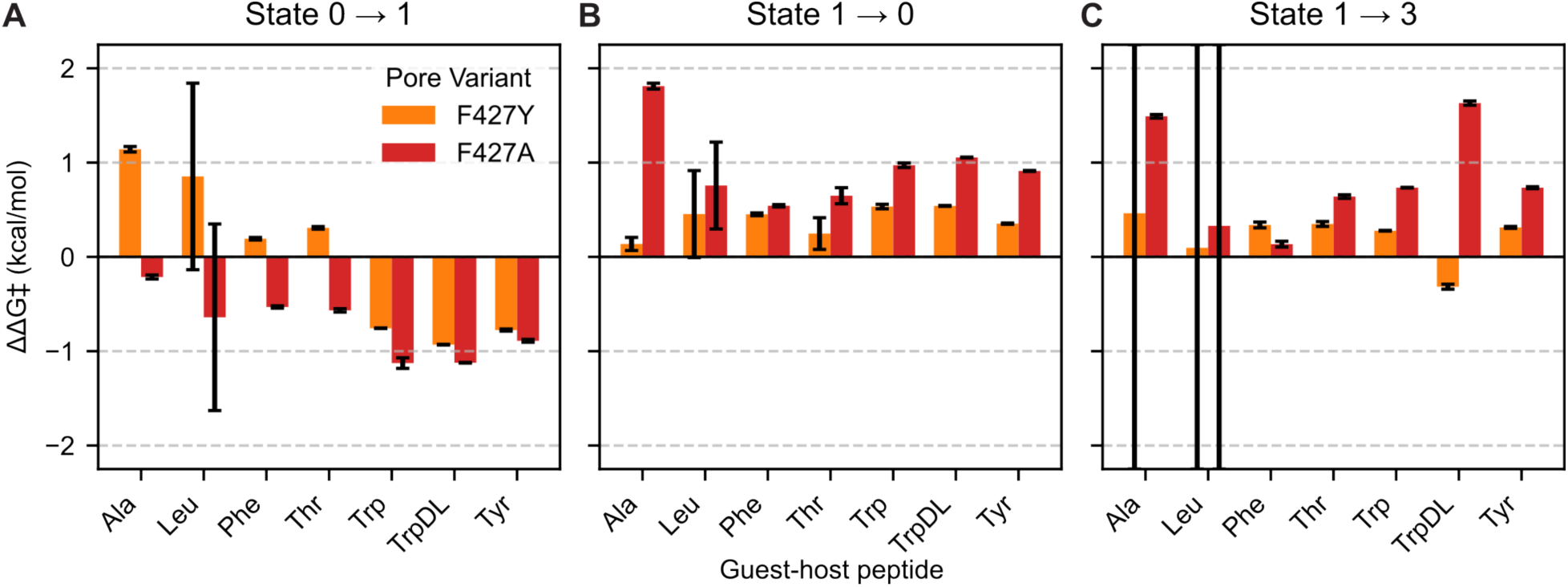
Kinetic analysis reveals the state-dependent energy barriers of translocation. The change in the activation free energy (ΔΔG‡) for key transitions in the PA F427A (red) and PA F427Y (orange) mutants relative to the wild-type PA nanopore for the indicated guest-host peptides (*x* axes). ΔΔG‡ was calculated from mean lifetimes as *RT* ln(τ_mutant_ / τ_WT_), where a negative value indicates the mutation lowered the energy barrier, making the transition faster. The representative panels shown are selected phases of the translocation mechanism. **(A)** The 0→1 transition represents the subsequent escape, a process governed by sterics and gated by aromaticity; its barrier is lowered by the F427A mutation. **(B)** The 1→0 transition represents the entry into the deep hydrophobic trap; its barrier is increased by F427A mutation. **(C)** The 1→3 transition represents the final dissociation from the intermediate state, a process limited by the strength of hydrophobic and aromatic interactions. The full set of nine kinetic transitions is shown in **Fig. S3**.

### Peptide hydrophobicity/aromaticity does not predict kinetics in F427A pores

From the consolidated amplitudes and the log of the lifetimes describing each transition a linear regression correlation analysis was performed, comparing those kinetic parameters to the molecular properties of the guest residues in the peptides. From the prior correlation analysis of wild-type PA, aromaticity metrics (aromatic residue Boolean and number of rings) and hydrophobicity scales based on residue transfer free energies (Hopp-Woods and Song) scored well (14). However, comparatively these correlations were much more muted and even entirely absent in the F427Y and F427A nanopores, respectively **(Fig. S4, Supporting Document 5)**. The implication of this result is that the F427 residue acts as a specific chemical reader that is highly sensitive to the hydrophobicity and aromaticity properties of the guest residue. When this reader is ablated (F427A), the interactions become generic and non-specific, erasing the predictive power of the simple physical properties. This serves as strong evidence that the correlations observed in the wild-type are not coincidental but are a direct consequence of specific interactions with the ϕ-clamp site.

## Discussion

### The ϕ clamp as a state-dependent dynamic gate

Our results demonstrate that the F427 residue is not a static interaction site but a key component of a dynamic gate that governs the multi-state translocation of peptides. By employing targeted mutations (F427A and F427Y), we have been able to systematically perturb the translocation energy landscape, providing an unprecedented high-resolution map of the ϕ-clamp’s function. This approach reveals a complex interplay of forces where the clamp acts not just as a physical constriction but as an active participant in the translocation process.

A central finding of this study is the role of the F427 residue as a specific chemical ‘reader’ that is highly sensitive to the molecular properties of the guest residue. In the wild-type pore, the kinetics of translocation show strong and predictable correlations with fundamental properties like side-chain hydrophobicity and aromaticity (14). However, these correlations are dramatically weakened in the F427Y mutant and are almost completely abolished upon ablation of the aromatic side chain in the F427A mutant **(Fig. S4, Supporting Document 5)**. This stark loss of predictive power demonstrates that the interactions within the wild-type clamp are not generic or merely steric; they are chemically specific, driven by the unique ability of the phenylalanine ring to form favorable hydrophobic and aromatic interactions. When this reader is removed, the peptide-pore interactions become non-specific, erasing the clear relationship between side-chain chemistry and translocation dynamics. This serves as powerful evidence that the correlations observed in the wild-type pore are a direct consequence of specific interactions with the F427 ϕ-clamp site.

### Refining the structural model: ϕ-clamp gating may encompass larger loop

The previously described model (14), where State 1 and, in particular, State 2 are peptide liganded at the ϕ clamp but are partially conducting states of the nanopore, is based primarily on guest-host translocation studies made via wild-type PA (**Fig. 4A**). It could be reasoned that the F427 phenyl moiety simply samples alternate rotamers in State 1 and State 2, creating room for potassium ions to permeate while a peptide is bound in the vicinity of the ϕ clamp. However, considering this new F427A variant nanopore data, the model can now be updated. This new finding is that intermediate, partially blocked conductance states (especially State 2) persist even in the F427A mutant, which lacks the aromatic side chain entirely (**Fig. 1C**). This finding rather supports a model where these sub-conductance states correspond to larger-scale conformational rearrangements—a ‘dilation’—of the entire clamp-containing loop (**Fig. 4B**). This establishes the loop itself as the primary gating element. Modeling the prepore heptamer ϕ-clamp loop (20) onto the heptameric pore shows that this potential natural loop conformation may lead to a dilated arrangement of the clamp (**Fig. 4B**). It should be noted that the solved structure of the ϕ-clamp loop in the heptameric pore matches the conformation of the clamp loop in the prepore PA octamer (2, 21); therefore, our putative model is within reason for this system.

**Fig. 4.**
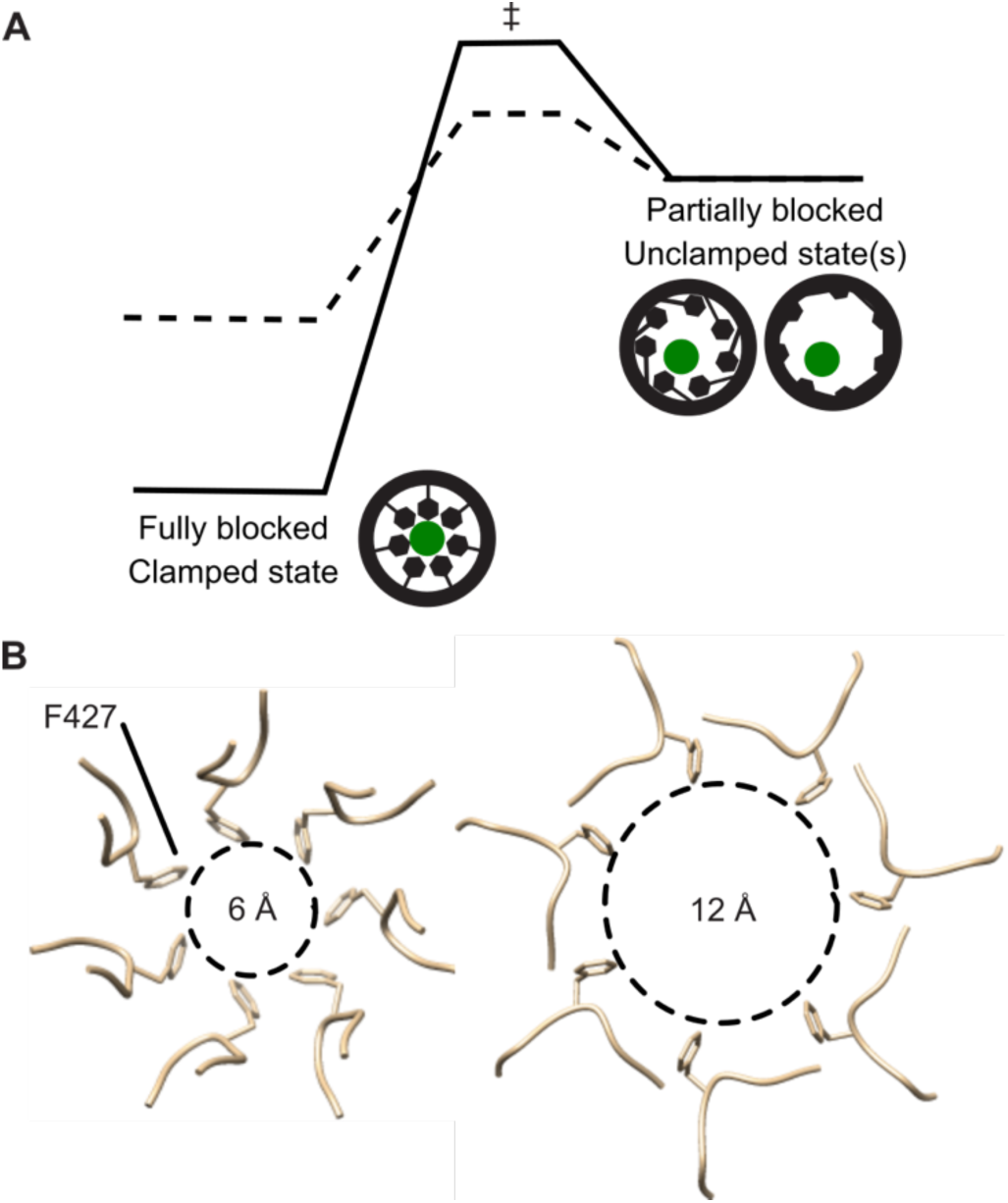
A refined model for dynamic peptide-clamp gating and its underlying energetics. **(A)** A schematic free energy landscape illustrates the key effects of the F427A mutation on the translocation process for a representative aromatic peptide. The solid line represents the wild-type pore, featuring a deep energy well corresponding to the stable, hydrophobic trap of State 0. The dashed line represents the F427A mutant, which significantly destabilizes State 0 (shallowing the well) and lowers the activation energy barrier (ΔG‡) for escape from this trapped state, consistent with the kinetic and thermodynamic data. Cartoons show ϕ-clamp (black) engagement with peptide (green) in constricted and various more dilated conformers. **(B)** (left) Cryo-EM structure (2) of the narrow, constricted ϕ-clamp loop in the PA heptamer, with the 6-Å luminal opening defined by the F427 residues (sticks). When liganded with peptide this loop conformation corresponds to the fully blocked State 0. (right) A putative model of the dilated conformation of the ϕ-clamp loop, corresponding to the partially conducting State 2 when liganded with peptide. This model, based on alternate loop conformations borrowed from the prepore heptamer (20), proposes a larger-scale rearrangement of the entire clamp loop is responsible for gating, rather than a simple rotamer change of the F427 phenyl residue.

Despite the suitability of PA as a model system, a critical discrepancy exists between structural and functional data regarding the channel’s ϕ-clamp site. On one hand, all current cryo-EM structures consistently show a static, constricted 6-Å luminal diameter opening (**Fig. 4B**) (2, 4, 5). This constriction point is sterically challenging for the passage of even single residues with large side chains and would completely occlude any transient helical secondary structure (22). However, a large body of functional electrophysiological evidence amassed over the past several decades presents a different picture, one of a dynamic and dilatable channel. Tetraalkylammonium sizing ions and diffusion studies suggest a much larger 11-12 Å diameter for the narrowest point of the channel (23, 24). More compellingly, single-channel measurements, albeit at near neutral pH, directly demonstrate the existence of alternate conductance states, suggesting the narrow opening can dynamically dilate to a larger diameter (9, 25). The dilation is also supported by studies showing a preformed, stapled helical peptide can translocate through the channel (11), which would be impossible via a 6-Å opening. Finally, single-channel peptide-binding studies show that the ϕ-clamp site exists in at least two different thermodynamic states under allosteric control, i.e., a tighter clamped state and a weaker unclamped state, and that the site is highly dynamic (9). Some researchers, nonetheless, remain skeptical and have argued against a dilated ϕ-clamp configuration, albeit those experiments neither used polypeptide substrates nor probed the partially conducting peptide-channel co-complexed intermediates directly (26).

### Dissecting the energy landscape: thermodynamics and kinetics

Based on our thermodynamic analysis, the F427 residue in the ϕ-clamp loop is a primary site of peptide interaction when forming the deep, stable energy well of State 0 (where ΔΔG > 0) (**Fig. 2, 4A)**. This is not surprising given earlier results showing a large loss of binding free energy change for tetrabutylammonium and tetraphenylphosphonium ions when comparing wild-type PA nanopore to the F427A variant (15). Hence, we have dubbed the formation of the deep State 0 peptide-clamp complex as a ‘hydrophobic trap’. The other consequences of destabilizing State 0 with F427A and F427Y mutations is that the system tends to populate State 1 and State 2 more significantly in those mutant contexts. This population shift to States 1 and 2 is a classic thermodynamic partitioning effect.

Kinetic results (**Fig. 3, S3)** tell an interesting story about the changes in barrier heights when traversing the energy landscape during a translocation event. The F427 residue plays a dual, state-dependent role: it creates a high activation barrier for escape from the trap (0→1) and for final dissociation (1→3), thus acting as the primary determinant of longer residence times than the mutant nanopores. The other escape route from the hydrophobic trap (0→2) is also more favorable in the F427X mutant backgrounds particularly for aromatic peptides. Paradoxically, F427 also facilitates the efficient, hydrophobically-driven entry into the trap (1→0), a process disrupted by the mutations. Thus, these kinetic results support the overall view of the energy landscape, where F427X mutants disrupt route into and out of the hydrophobic trap State 0.

### Implications for translocation and peptide biosensing

From these results and previous studies on the role of the ϕ-clamp site, an interesting translocase vs. sensor paradox is emerging (**Table 1**). PA F427A is a very defective translocase compared to wild-type or F427Y for larger folded proteins in cellular assays and ensemble electrophysiology studies, especially under proton gradient driving forces (15). However, F427A is the most capable biosensor and classifier for these small guest-host peptides with up to 93% accuracy (13). The strong dependence of the multi-state translocation kinetics on hydrophobicity and aromaticity in the guest residue positions for the wild-type PA system (14) **(Fig. S4)** is not requisite for strong biosensing classification performance (13). We can partly explain this phenomenon with a ‘low signal, low variance’ model. The wild-type F427-containing clamp produces a strong, specific signal but with high kinetic variance. The ablated F427A clamp produces a weaker, more generic signal, but the resulting kinetic ‘fingerprints’ are more consistent and less variable, making them easier for a machine learning classifier to distinguish.

**Table 1.**
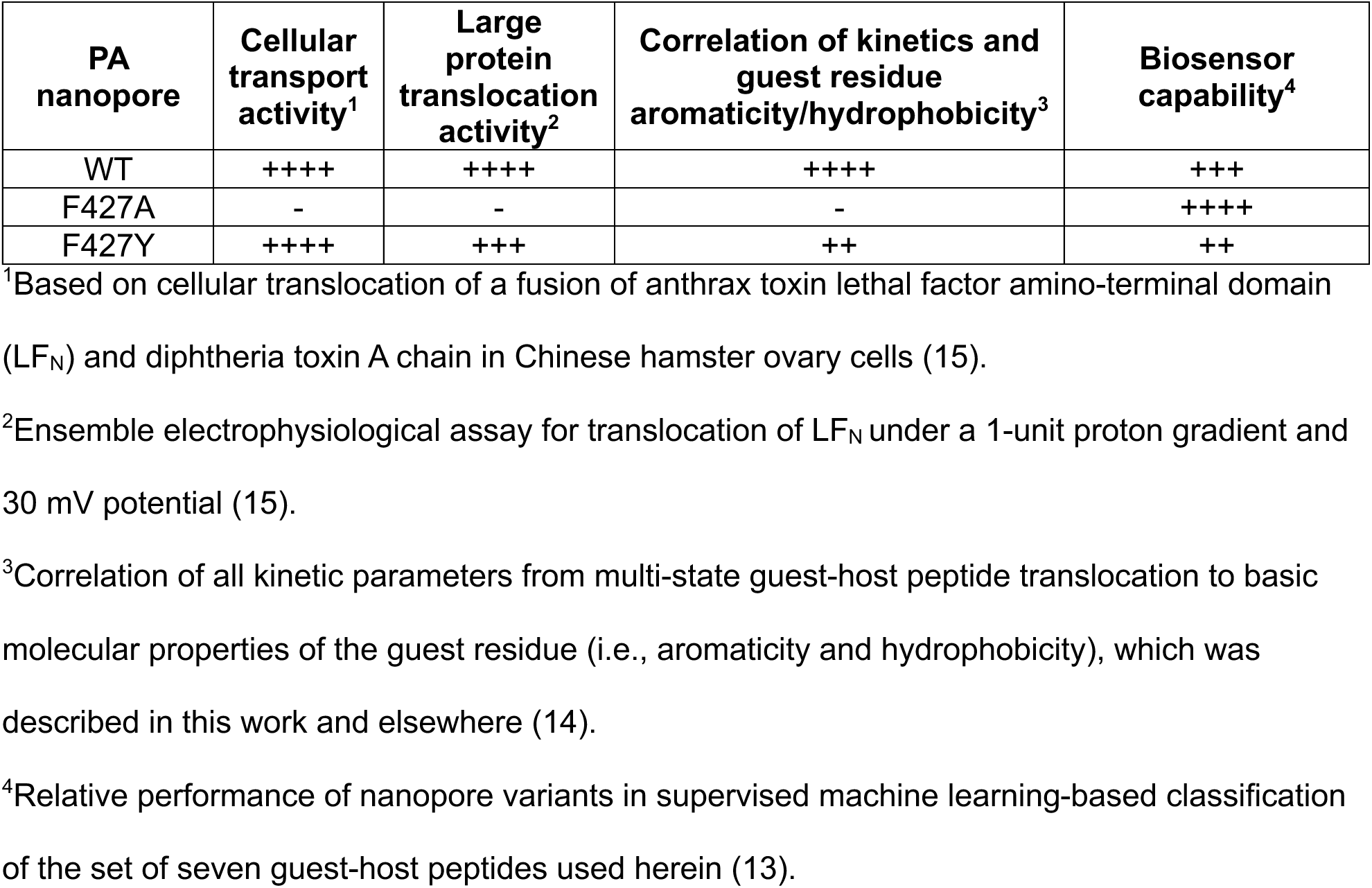
Relative performance of PA F427X variants in translocation and biosensing.

### Future Directions

Based on these insights, future work should pursue the structure, mechanism, and biosensing capabilities of this nanopore. While challenging potentially to produce high enough proportions of dilated conformers, structural approaches like cryo-EM can elucidate atomic resolution models even in mixed populations. Molecular dynamics simulations may be able to visualize our proposed loop dilation, or even folding prediction approaches may be able to discover alternate loop configurations. Mutagenesis and protein engineering may further probe other residues in the ϕ-clamp loop to determine the relationship between its confirmational dynamics and the overall translocation mechanism. These mechanistic insights and newly developed nanopore variants may be used to further rationally engineer the PA nanopore for enhanced biosensing and even sequencing applications.

## Conclusion

In many soluble and membrane embedded translocases ϕ-clamp-like active sites play a prominent functional role. The work described here provides a comprehensive dissection of the ϕ clamp’s function in the mechanism of peptide translocation. Far from the earlier models, the clamp is not likely a static structure with a fixed 6-Å luminal diameter, but it is instead a dynamic, multi-state gate that significantly dilates. Its properties arise from both the specific chemistry of the F427 residue and the conformational flexibility of the entire loop. This dual nature explains its seemingly paradoxical roles as both an efficient protein unfoldase/translocase and a high-fidelity peptide biosensor.

## Materials and Methods

### Nanopore guest-host peptide system and data acquisition

The wet lab procedures and the collection of raw datasets used in this analysis were previously described (12, 13). Briefly, monomeric 83-kDa PA, PA F427A and PA F427Y preprotein and their homoheptameric prepore oligomers (PA_7_) were produced as described (15, 21). Ten-residue guest-host peptides of the general sequence, KKKKKXXSXX, where X = A, L, F, T, W, and Y, were synthesized with standard ʟ amino acids (8, 11–13, 27) (Elim Biopharmaceuticals). One stereochemical variant of X = W (called TrpDL) was produced, where instead of synthesizing the peptide with uniform ʟ amino acids, an alternating pattern of ᴅ and ʟ amino acids was used.

Planar lipid bilayers were used to acquire peptide translocation event streams via PA nanopores and variants using an Axopatch 200B amplifier system (Molecular Devices) as previously described (11, 12, 21, 28). The basic experimental conditions used in peptide translocation recordings were 100 mM KCl, 20 mM succinic acid, 1 mM EDTA, pH 5.6 under a constant +70 mV driving force (cis positive). A 400 Hz sampled dataset was used, where in some cases higher time sampled raw data were downsampled to 400 Hz. Conductance states were labeled using a computationally efficient K-Means clustering routine (13). By convention, the fully blocked peptide-bound state was State 0, the intermediate closest to the fully blocked state was State 1, the intermediate closest to the open state was State 2, and the open state was State 3. All labeled raw datasets were entered into a local database to aid in processing large numbers of translocation event streams. Centroids from K-Means labeling of conductance states were used to determine the fractional conductance blockades of intermediates, State 1 and State 2, for all raw streams for all guest-host peptides for each nanopore variant. Centroids for the dynamic range of the nanopore variants were used to determine the conductance of the fully open nanopores.

### Translocation event segmentation

Raw state-labeled event streams were segmented into translocation events, as described (12). The minimum event duration, which served as an effective filter for excluding very short events which can be unrelated to peptide translocation events, was set to 5 ms to ignore the shortest 2.5 ms wide spike events (single time point at 400 Hz sampling). Each segmented event was defined as initiating when the current changed from the fully open state (State 3, corresponding to baseline current) to any peptide-bound state (State 0, 1, or 2) and terminating when the current returned to State 3. From these segmented events, the corresponding state sequences were extracted. Dwell times were computed for each transition from State *i* to State *j* creating a 4×4 transition matrix.

### Thermodynamic analysis

From the dwell times in transition matrix, the residence time in each state encountered in a translocation event (State 0, 1, and 2) were computed to then obtain the probability in each state, *P*_i_. The change in free energy for each nanopore variant for each tested guest-host peptide was calculated

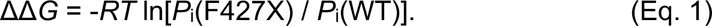

### Kinetic analysis

A cumulative distribution function (CDF), survival curve, *S* (1 - CDF), and natural log of the survival curve, ln(*S*), were determined for each position in the dwell time transition matrix. Different exponential decay models were fitted to ln(*S*), including single-(Eq. 2), double-(Eq. 3), and triple-exponential decay functions (Eq. 4), yielding respective lifetimes, τ, and amplitudes, *A*.

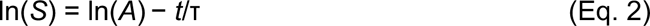

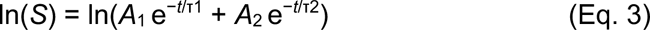

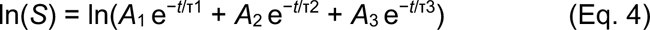

To enable comparison of complex multi-exponential kinetic transitions to single-exponential ones, mean lifetime (τ_mean_) was calculated by τ_mean_ = Σ τ_i_*A*_i_. The best exponential model for each transition for each peptide for each nanopore was determined by the Bayesian Information Criterion (BIC), which penalizes over parameterization. In terms of the residual sum of squares (RSS) from the respective curve fits, BIC = *n* ln(RSS/*n*) + *k* ln(*n*). The best exponential decay model was then selected based on the lowest BIC, ignoring, of course, all fits that failed to converge. To aid in downstream comparative analysis the lifetimes for different exponential decay models were handled systematically by ranking as τ_fast_, τ_middle_, and τ_slow_. When the model was a one-exponential decay, then only τ_fast_ was used. When a two-exponential model was used τ_fast_ was the fastest decay, and τ_middle_ and τ_slow_ were the same second, slower decay lifetime. The respective fast, middle and slow amplitudes were handled similarly as the τ values in this consolidation process.

Estimates of energy barrier heights between the observed transitions (Δ*G*‡) were determined from τ_mean_ values by Δ*G*‡ = *RT* ln τ_mean_. The change in the activation free energy (ΔΔ*G*‡) was calculated to quantify how the F427X mutations affect the energy barriers for specific transitions between states: ΔΔ*G*‡ = Δ*G*‡(F427X) - Δ*G*‡(WT).

### Correlations of kinetic parameters to guest residue molecular properties

The consolidated kinetic parameter data for the observed transitions for each peptide were then analyzed by correlating the molecular properties of the guest residue to the base-10 log(τ) and raw *A* values, including τ_mean_, τ_fast_, τ_middle_, τ_slow_, *A*_fast_, *A*_middle_, and *A*_slow_, according to a previously described procedure (14). The molecular properties of the guest residue included: molecular weight; an aromaticity Boolean; number of aromatic rings; various hydrophobicity scales (Kyte-Doolittle (29), Hopp-Woods (30), Cornette (31), Eisenberg (32), Rose (33), Janin (34), Engelman GES (35), Tanford (36), and Song (37)); and two atomic solvation parameterized energy scales (Ooi (38) and Krantz (15)). From these scales, the molecular property data for the 20 natural amino acids were compiled **(Supporting Document 6)**. Linear regression analysis was performed between the kinetic parameters and molecular properties for each transition observed to generate *R*^2^ values for each correlation **(Supporting Document 5)**. The best overall molecular properties (Mol. wt., Aromaticity, Num. rings, Hopp-Woods, and Song), which were most predictive for wild-type PA over all kinetic parameters and transitions, were assessed for their correlative ability in mutant nanopores by mean *R*^2^ values over all parameters for all observed kinetic transitions.

## Data availability statement

All experimental electrophysiological records (with K-Means state labeling) and related source code are publicly available. The datasets used in this manuscript have been deposited in the Zenodo repository under the DOI: 10.5281/zenodo.17203719. The source code is maintained on a GitHub repository (https://github.com/bakrantz/Pept-Class).

## Acknowledgments

We thank members of the department for useful feedback and discussions. J.M.C. and B.A.K. conceived of the experiments. J.M.C. collected the data. J.M.C. and B.A.K performed analyses. B.A.K. and J.M.C. wrote the manuscript. Portions of this document, including some of the Python code and language refinement, were generated with the assistance of AI-powered tools. All content was reviewed and approved by the authors, who take full responsibility for its accuracy.

---

**Table S1.**
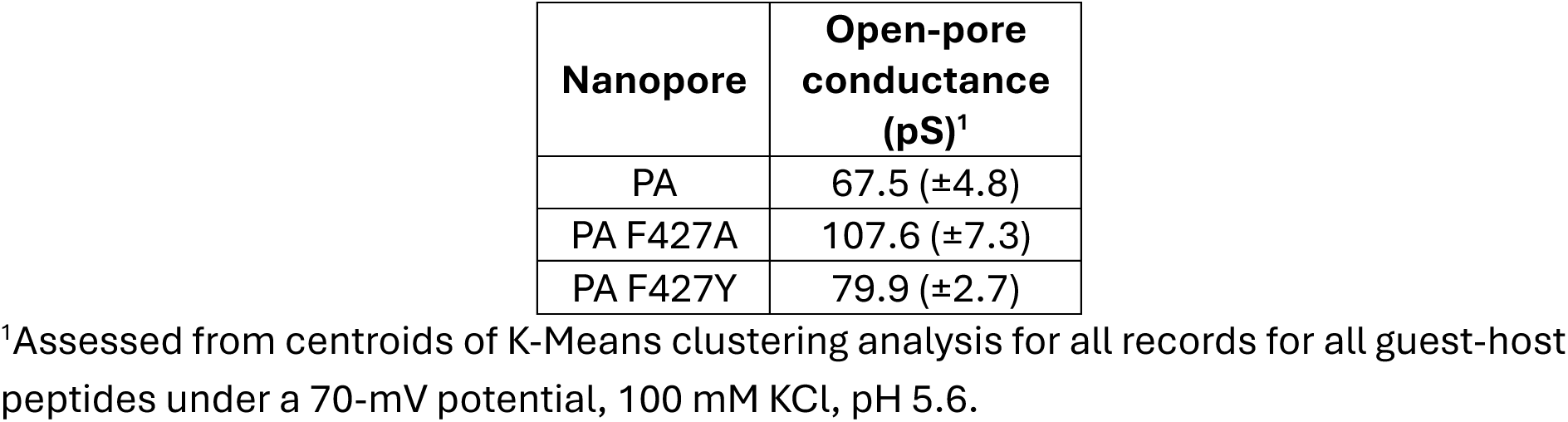
Open-pore conductance of wild-type PA and its ϕ-clamp variants.

**Fig. S1.**
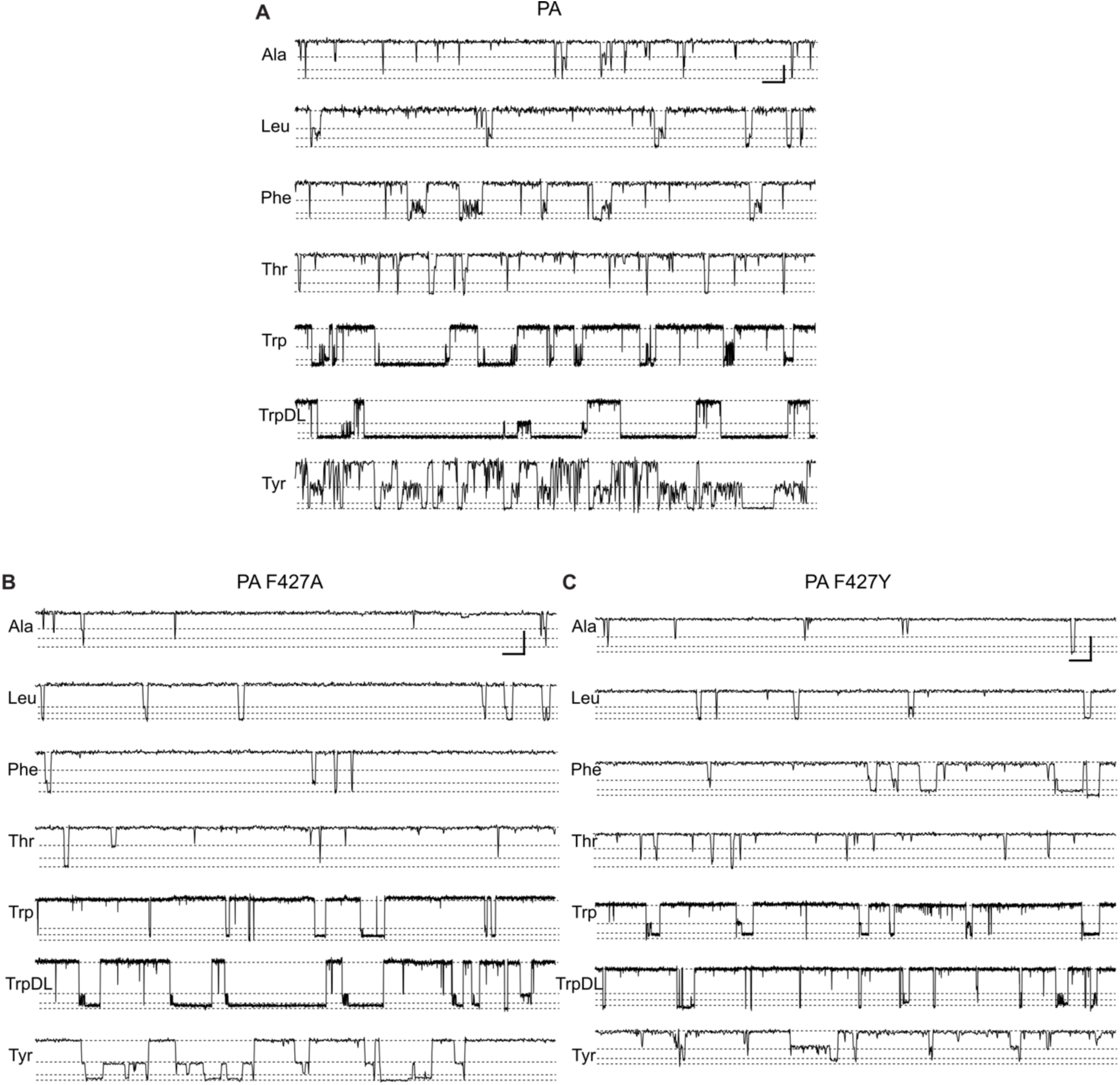
Samples of translocation event streams for guest-host peptides via variant PA nanopores. Representative samples of recordings of guest-host peptide (20 nM) translocation event streams via **(A)** wild-type PA, **(B)** PA F427A, and **(C)** PA F427Y nanopores, which taken at 70 mV (cis positive) in symmetric 100 mM KCl, 20 mM succinate, 1 mM EDTA, pH 5.6. To the left are the standard three letter name for the guest residue. These nanopore-peptide systems populate multiple discrete partially or fully blocked intermediates (approximate levels indicated by dotted lines). From bottom to top of each record: fully blocked (State 0), partially blocked intermediates (State 1 and State 2), and fully open baseline (State 3). Scalebar at the upper right of panel A denotes 2 pA by 100 ms for guest-host Ala, Leu, Phe, Thr, and Tyr peptides, but for guest-host Trp and TrpDL peptides, the scalebar is 2 pA by 500 ms to present longer events. Scalebar at the upper right of panels B and C denotes 4 pA by 100 ms for guest-host Ala, Leu, Phe, Thr, and Tyr peptides, but for guest-host Trp and TrpDL peptides, the scalebar is 4 pA by 500 ms to present longer events.

**Fig. S2.**
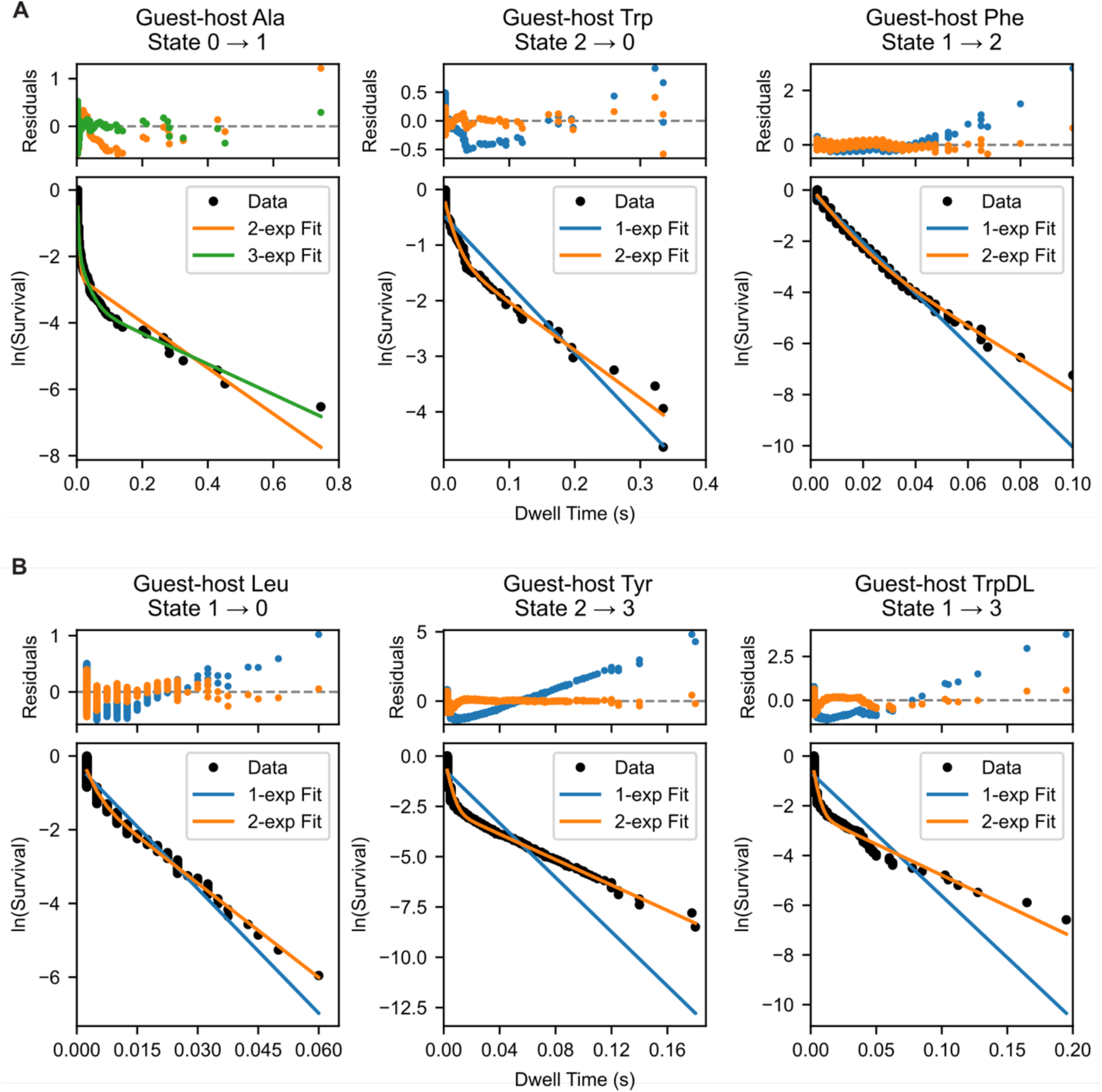
Translocation event kinetics generally fit to multi-exponential models. Natural log of the survival curves plotted against dwell time (black circles) are shown for selected translocation event transitions for guest-host peptides via **(A)** PA F427A and **(B)** PA F427Y nanopores. Exponential decay models, colored by number of exponentials (1-exp, blue; 1-exp, orange; 2-exp, green), were selected using BIC. The residuals plot at top of each panel, colored consistently with the exponential fit models, further justified these model selections.

**Fig. S3.**
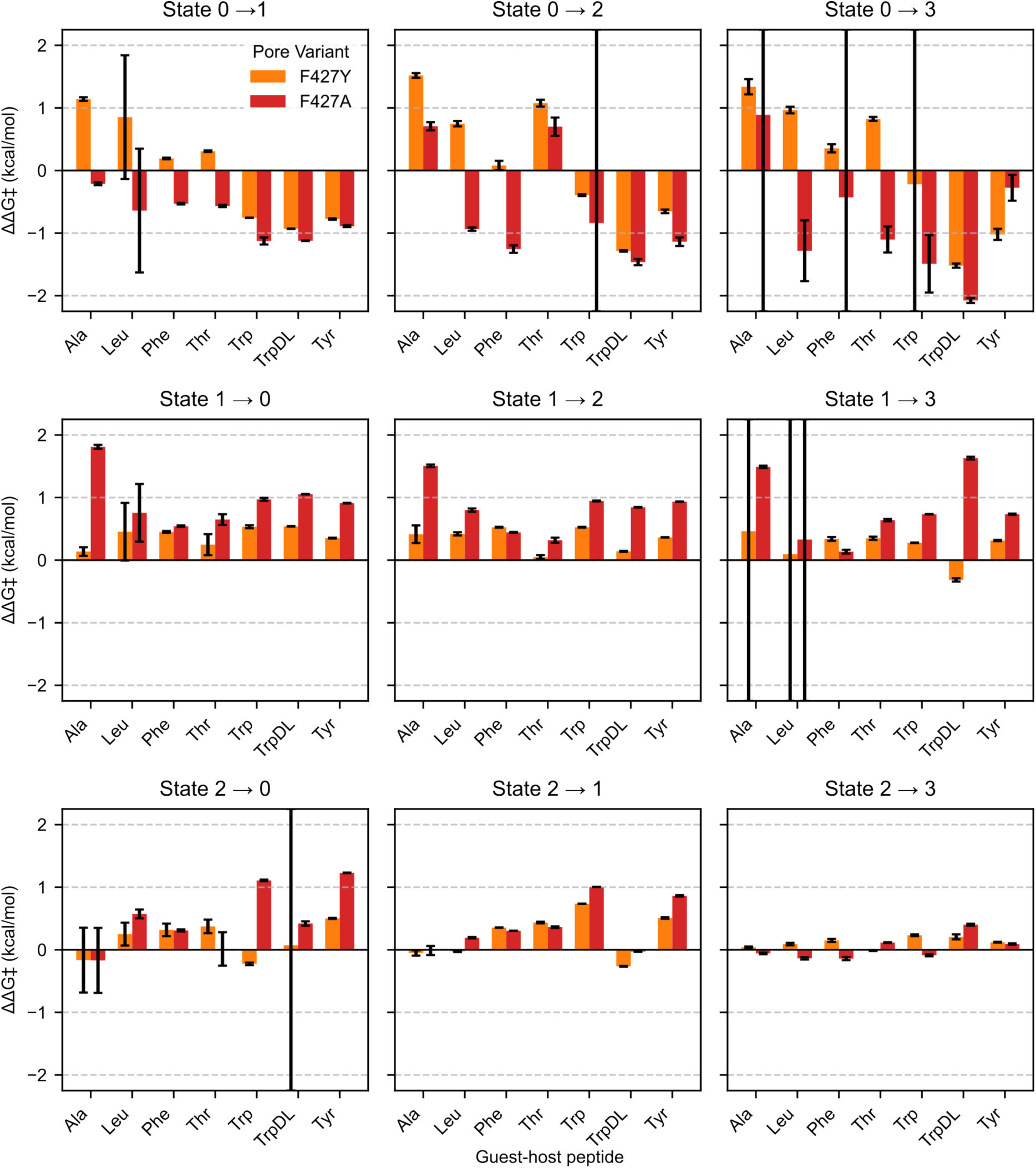
Comprehensive kinetic analysis of peptide-clamp interactions in all observed state transitions. The change in the activation free energy (ΔΔG‡) for all nine observed forward transitions in the PA F427A (red bars) and PA F427Y (orange bars) mutants relative to the wild-type pore. ΔΔG‡ was calculated from mean lifetimes as *RT* ln(τ_mutant_ / τ_WT_); negative values indicate the mutation lowered the energy barrier, making the transition faster, while positive values indicate a higher barrier. The data reveal a complex, state-dependent response to the mutations. Note the significant lowering of the energy barriers (negative ΔΔG‡) for transitions originating from the fully blocked State 0, particularly for the F427A mutant with aromatic guest residues.

**Fig. S4.**
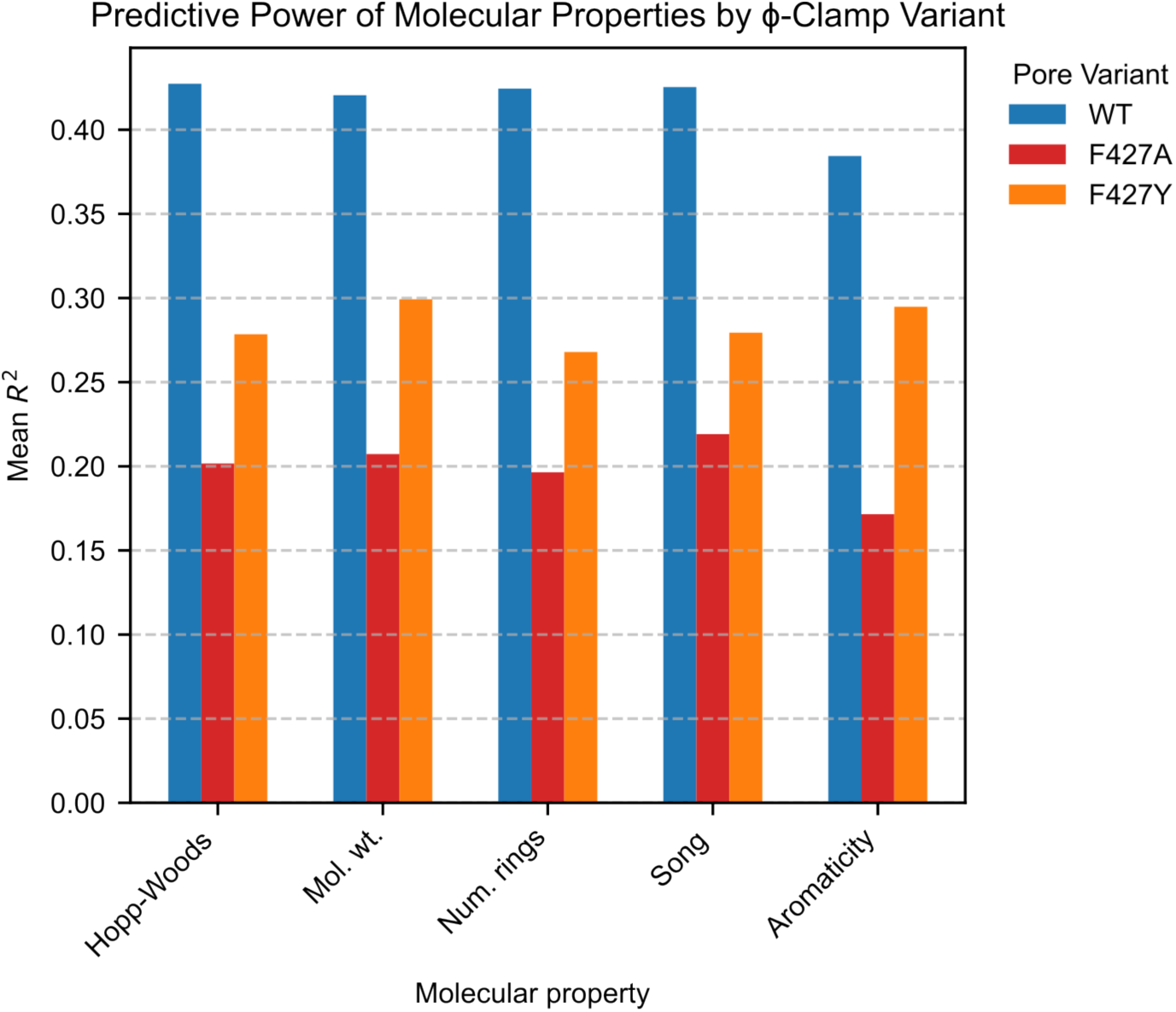
Correlations of molecular properties and peptide translocation kinetic parameters for ϕ variants. Mean *R*^2^ correlations of linear regressions between translocation event kinetic parameters for all observed transitions versus selected molecular properties of the guest residue in the peptides. Correlations for wild-type PA (blue), PA F427A (red), and PA F427Y (orange). Selected properties shown here are: a hydrophobicity scale (Hopp-Woods) (30), molecular weight (Mol. wt.), number of aromatic rings (Num. rings), a transfer free energy scale (Song) (37), and Boolean for whether the guest residue is aromatic (Aromaticity). Comprehensive correlations between all tested molecular properties with a breakdown for each kinetic parameter are also available **(Supporting Document 5)**.

## References

1. B. A. Krantz, Anthrax Toxin: Model System for Studying Protein Translocation. J Mol Biol 436, 168521 (2024).

2. J. Jiang, B. L. Pentelute, R. J. Collier, Z. H. Zhou, Atomic structure of anthrax protective antigen pore elucidates toxin translocation. Nature 521, 545–549 (2015).

3. G. K. Feld et al., Structural basis for the unfolding of anthrax lethal factor by protective antigen oligomers. *Nature Struct*. Mol. Biol. 17, 1383–1390 (2010).

4. N. J. Hardenbrook et al., Atomic structures of anthrax toxin protective antigen channels bound to partially unfolded lethal and edema factors. Nat Commun 11, 840 (2020).

5. A. J. Machen, M. T. Fisher, B. D. Freudenthal, Anthrax toxin translocation complex reveals insight into the lethal factor unfolding and refolding mechanism. Sci Rep 11, 13038 (2021).

6. S. Zhang, E. Udho, Z. Wu, R. J. Collier, A. Finkelstein, Protein translocation through anthrax toxin channels formed in planar lipid bilayers. Biophys. J. 87, 3842–3849 (2004).

7. B. A. Krantz, A. Finkelstein, R. J. Collier, Protein translocation through the anthrax toxin transmembrane pore is driven by a proton gradient. J. Mol. Biol. 355, 968–979 (2006).

8. J. M. Colby, B. A. Krantz, Peptide Probes Reveal a Hydrophobic Steric Ratchet in the Anthrax Toxin Protective Antigen Translocase. J Mol Biol 427, 3598–3606 (2015).

9. D. Das, B. A. Krantz, Peptide- and proton-driven allosteric clamps catalyze anthrax toxin translocation across membranes. Proc Natl Acad Sci U S A 113, 9611–9616 (2016).

10. D. Das, B. A. Krantz, Secondary Structure Preferences of the Anthrax Toxin Protective Antigen Translocase. J Mol Biol 429, 753–762 (2017).

11. K. Ghosal et al., Dynamic Phenylalanine Clamp Interactions Define Single-Channel Polypeptide Translocation through the Anthrax Toxin Protective Antigen Channel. J Mol Biol 429, 900–910 (2017).

12. J. M. Colby, B. A. Krantz, High-performance machine learning for peptide classification from nanopore translocation events, leveraging event kinetics and duration filtering. bioRxiv 10.1101/2025.08.21.671556 (2025).

13. J. M. Colby, B. A. Krantz, Comparative study of nanopore phenylalanine clamp variants reveals unique peptide biosensing and classification properties. bioRxiv 10.1101/2025.08.22.671566 (2025).

14. J. M. Colby, B. A. Krantz, Peptide hydrophobicity and aromaticity predict multi-state translocation kinetics via protective antigen nanopores. bioRxiv 10.1101/2025.09.15.676337 (2025).

15. B. A. Krantz et al., A phenylalanine clamp catalyzes protein translocation through the anthrax toxin pore. Science 309, 777–781 (2005).

16. S. L. Wynia-Smith, M. J. Brown, G. Chirichella, G. Kemalyan, B. A. Krantz, Electrostatic ratchet in the protective antigen channel promotes anthrax toxin translocation. J Biol Chem 287, 43753–43764 (2012).

17. B. Lin, J. Hui, H. Mao, Nanopore Technology and Its Applications in Gene Sequencing. Biosensors (Basel) 11 (2021).

18. Y. Goto, R. Akahori, I. Yanagi, Challenges of Single-Molecule DNA Sequencing with Solid-State Nanopores. Adv Exp Med Biol 1129, 131–142 (2019).

19. X. Wei et al., Engineering Biological Nanopore Approaches toward Protein Sequencing. ACS Nano 17, 16369–16395 (2023).

20. D. B. Lacy, D. J. Wigelsworth, R. A. Melnyk, S. C. Harrison, R. J. Collier, Structure of heptameric protective antigen bound to an anthrax toxin receptor: a role for receptor in pH-dependent pore formation. Proc. Natl. Acad. Sci. U.S.A. 101, 13147–13151 (2004).

21. A. F. Kintzer et al., The protective antigen component of anthrax toxin forms functional octameric complexes. J. Mol. Biol. 392, 614–629 (2009).

22. S. Gonti, W. M. Westler, M. Miyagi, J. G. Bann, Site-Specific Labeling and (19)F NMR Provide Direct Evidence for Dynamic Behavior of the Anthrax Toxin Pore varphi-Clamp Structure. Biochemistry 60, 643–647 (2021).

23. R. O. Blaustein, A. Finkelstein, Voltage-dependent block of anthrax toxin channels in planar phospholipid bilayer membranes by symmetric tetraalkylammonium ions. Effects on macroscopic conductance. J Gen Physiol 96, 905–919 (1990).

24. R. O. Blaustein, A. Finkelstein, Diffusion limitation in the block by symmetric tetraalkylammonium ions of anthrax toxin channels in planar phospholipid bilayer membranes. J Gen Physiol 96, 943–957 (1990).

25. R. O. Blaustein, E. J. Lea, A. Finkelstein, Voltage-dependent block of anthrax toxin channels in planar phospholipid bilayer membranes by symmetric tetraalkylammonium ions. Single-channel analysis. J. Gen. Physiol. 96, 921–942 (1990).

26. N. Kalu et al., Effect of endosomal acidification on small ion transport through the anthrax toxin PA(63) channel. FEBS Lett 591, 3481–3492 (2017).

27. B. A. Krantz, Deep learning-based classification of peptide analytes from single-channel nanopore translocation events. PLoS One 20, e0324777 (2025).

28. K. L. Thoren, E. J. Worden, J. M. Yassif, B. A. Krantz, Lethal factor unfolding is the most force-dependent step of anthrax toxin translocation. Proc. Natl Acad. Sci. U.S.A. 106, 21555–21560 (2009).

29. J. Kyte, R. F. Doolittle, A simple method for displaying the hydropathic character of a protein. J Mol Biol 157, 105–132 (1982).

30. T. P. Hopp, K. R. Woods, A computer program for predicting protein antigenic determinants. Mol Immunol 20, 483–489 (1983).

31. J. L. Cornette et al., Hydrophobicity scales and computational techniques for detecting amphipathic structures in proteins. J Mol Biol 195, 659–685 (1987).

32. D. Eisenberg, E. Schwarz, M. Komaromy, R. Wall, Analysis of membrane and surface protein sequences with the hydrophobic moment plot. J Mol Biol 179, 125–142 (1984).

33. G. D. Rose, A. R. Geselowitz, G. J. Lesser, R. H. Lee, M. H. Zehfus, Hydrophobicity of amino acid residues in globular proteins. Science 229, 834–838 (1985).

34. J. Janin, Surface and inside volumes in globular proteins. Nature 277, 491–492 (1979).

35. D. M. Engelman, T. A. Steitz, A. Goldman, Identifying nonpolar transbilayer helices in amino acid sequences of membrane proteins. Annu Rev Biophys Biophys Chem 15, 321–353 (1986).

36. Y. Nozaki, C. Tanford, The solubility of amino acids and two glycine peptides in aqueous ethanol and dioxane solutions. Establishment of a hydrophobicity scale. J Biol Chem 246, 2211–2217 (1971).

37. S. Damodaran, K. B. Song, The role of solvent polarity in the free energy of transfer of amino acid side chains from water to organic solvents. J Biol Chem 261, 7220–7222 (1986).

38. T. Ooi, M. Oobatake, G. Nemethy, H. A. Scheraga, Accessible surface areas as a measure of the thermodynamic parameters of hydration of peptides. Proc Natl Acad Sci U S A 84, 3086–3090 (1987).

